# Mitochondrial choline import regulates purine nucleotide pools via SLC25A48

**DOI:** 10.1101/2023.12.31.573776

**Authors:** Anthony R.P. Verkerke, Xu Shi, Ichitaro Abe, Robert E. Gerszten, Shingo Kajimura

**Affiliations:** Division of Endocrinology, Diabetes and Metabolism, Beth Israel Deaconess Medical Center and Harvard Medical School, and Howard Hughes Medical Institute, Boston, MA, USA; Division of Cardiovascular Medicine, Beth Israel Deaconess Medical Center and Harvard Medical School, Boston, MA, USA; Department of Cardiology and Clinical Examination, Oita University, Faculty of Medicine, Oita, Japan

**Keywords:** Brown adipose tissue, Choline, Mitochondria, Cell viability, Cancer metabolism

## Abstract

Choline is an essential nutrient for cellular metabolism, including the biosynthesis of phospholipids, neurotransmitters, and one-carbon metabolism. A critical step of choline catabolism is the mitochondrial import and synthesis of chorine-derived methyl donors, such as betaine. However, the underlying mechanisms and the biological significance of mitochondrial choline catabolism remain insufficiently understood. Here, we report that a mitochondrial inner-membrane protein SLC25A48 controls mitochondrial choline transport and catabolism *in vivo*. We demonstrate that SLC25A48 is highly expressed in brown adipose tissue and required for whole-body cold tolerance, thermogenesis, and mitochondrial respiration. Mechanistically, choline uptake into the mitochondrial matrix via SLC25A48 facilitates betaine synthesis and one-carbon metabolism. Importantly, cells lacking SLC25A48 exhibited reduced synthesis of purine nucleotides and failed to initiate the G1-to-S phase transition, thereby leading to cell death. Taken together, the present study identified SLC25A48 as a mitochondrial carrier that mediates choline import and plays a critical role in mitochondrial respiratory capacity, purine nucleotide synthesis, and cell survival.

**Key points:**

- SLC25A48 is required for mitochondrial choline uptake.
- Mitochondrial choline uptake regulates one-carbon contribution to purine nucleotide synthesis.
- Brown fat thermogenesis requires mitochondrial choline catabolism for respiratory capacity.
- Cancer cells require mitochondrial choline uptake for cell survival.

## INTRODUCTION

While the apparent role of mitochondria is producing energy currency in the form of adenosine triphosphate (ATP), mitochondria also play crucial and diverse roles in catabolizing, synthesizing, and regulating various metabolic processes through compartmentalized metabolism. Mitochondria produce their own phospholipids that are biophysically favorable for the ultrastructure of tightly compacted cristae (Heden et al., 2016; Shiao et al., 1995). They also breakdown and generate cellular amino acids within the mitochondrial compartment (Schuler et al., 2021; Spinelli and Haigis, 2018). Through these processes, mitochondria produce essential metabolic intermediates and signaling molecules that immensely impact the nuclear transcriptional program and cellular function (Chakrabarty and Chandel, 2021; Kusminski et al., 2012; Lin et al., 2022). It is important to note that the import and export of mitochondria-compartmentalized metabolites is tightly regulated by the inner mitochondrial membrane (IMM). The IMM is impermeable to metabolites, and thus, the transport across the IMM relies on designated membrane-bound carrier proteins. For instance, pyruvate and serine are imported into the matrix through the mitochondrial pyruvate carriers (MPC1/2) and sideroflexin 1 (SFXN1), respectively (Bricker et al., 2012; Herzig et al., 2012; Kory et al., 2018). Among mitochondrial metabolite transport proteins, Solute Carrier Family 25 A (SLC25A) proteins belong to the largest family of carrier proteins that are mostly localized to the IMM (Palmieri, 2013; Ruprecht and Kunji, 2020). However, many of these proteins are considered “orphan” carriers as the specific substrates and biological functions remain uncharacterized. This leaves many unknowns regarding how metabolites enter or exit the mitochondrial matrix.

Choline is one of the metabolites whose catabolism depends on mitochondrial import and compartmentalization. It is a key nutrient that contributes to the synthesis of phospholipids and neurotransmitters, and is also a major one-carbon donor within the cell (Zeisel, 2006). The conversion of choline to its one-carbon donor betaine (trimethylglycine) occurs within the mitochondria where a mitochondria-localized enzyme choline dehydrogenase (CHDH) converts choline to the intermediate betaine aldehyde, which is then converted to betaine by aldehyde dehydrogenase 7 family member A1 (ALDH7A1) (Ueland, 2011). Betaine, derived from choline within the mitochondria, is exported to the cytosolic compartment where it serves as a methyl donor for methionine, contributing to the folate cycle. The dynamic exchange of choline and betaine across the mitochondrial membrane is mediated by the IMM-bound carrier proteins; however, the underlying mechanism and pathophysiology are not yet fully understood.

The functional characterization of mitochondrial carriers has been challenging, in part, due to technical limitations. For example, bioinformatic predictions of SLC25A proteins based on the sequence homology led to inconsistent results. Mammalian uncoupling protein 1 (UCP1, also known as SLC25A7) imports H^+^ into the mitochondrial matrix (Jones et al., 2023; Kang and Chen, 2023), whereas UCP2, which has high homology to UCP1, exchanges C4 metabolites (Vozza et al., 2014). Besides, some of these proteins are expressed in a cell/tissue-selective manner, and thus, cell type is a critical element in understanding the biological significance. Therefore, it is important to characterize mitochondrial carrier proteins using biologically relevant cells and *in vivo*. In this regard, brown fat serves as a useful discovery platform due to its highly enriched mitochondria where most, if not all, IMM proteins are abundantly expressed. In addition, brown adipocytes exhibit unique and robust features, including dense cristae structure, mitochondrial uncoupling respiration, and thermogenesis (Cohen and Kajimura, 2021). By employing the cellular tools combined with metabolomics and *in vivo* physiology, this study characterized the function of a previously unknown SLC25A protein, SLC25A48, as a determinant of mitochondrial choline catabolism.

## RESULTS

### SLC25A48 is required for optimal brown adipose tissue thermogenesis

We started with unbiased proteomics to identify mitochondrial proteins in brown adipose tissue (BAT) that are regulated in response to diet-induced obesity. To this end, we performed quantitative tandem mass tag proteomics in isolated mitochondria from the BAT of wild-type male mice fed standard diet or high-fat diet (60% kcal from fat) for 8 weeks. The analysis identified several mitochondrial proteins down-regulated by high-fat diet feeding. These proteins include the citrate carrier (SLC25A1), the dicarboxylate acid carrier (SLC25A10), a glutamate carrier (SLC25A22), and a carrier associated with Graves’ disease (SLC25A16). In contrast, there was only one SLC25A protein that be significantly upregulated in response to high-fat diet feeding, SLC25A48, of which the function has previously been unreported (**Figure 1A**). SLC25A48 is highly expressed in BAT in addition to the liver and kidney, while its expression is nearly undetectable in the heart and soleus muscle even though these tissues contain high levels of mitochondria (**Supplemental Fig. 1A**). While many SLC25A family members are predicted to localize to the inner mitochondrial member (IMM), several family members do not. For instance, SLC25A17 localizes to the peroxisome, while SLC25A46 and SLC25A50 localize to the outer mitochondrial member (OMM) (Abrams et al., 2015; Agrimi et al., 2012; Zaltsman et al., 2010). Accordingly, we performed a proteinase degradation assay in isolated mitochondria to determine whether SLC25A48 localizes to the IMM or OMM. In brown adipocytes expressing control empty-vector or SLC25A48 with Flag-tag (SLC25A48-Flag), we utilized differential centrifugation to enrich the mitochondrial fraction, which was subjected to treatment with proteinase K. In proteinase K treated mitochondria, OMM marker translocase of outer mitochondrial marker 20 (TOMM20) was degraded, while IMM marker ATP synthase F1 subunit alpha (ATP5A) was not disrupted, representing that the IMM was intact and not exposed to the proteinase K. SLC25A48 expression was present in the mitochondrial fraction and was not degraded by proteinase K treatment, demonstrating that SLC25A48 is localized to the inner mitochondrial membrane (**Figure 1B**). With this information in hand, we next examined whether SLC25A48 expression was regulated during adipogenesis in wild-type brown adipocytes. Indeed, over the course of brown adipocyte differentiation, SLC25A48 expression was rapidly induced (**Figure 1C**). Of note, this induction is not a general trend in mitochondrial carrier proteins; some SLC25A proteins, such as SLC25A4, were unchanged during brown adipogenesis (**Supplemental Fig. 1B**). Together, these data suggest the functional role of SLC25A48 in brown adipocytes.

**Figure 1.**
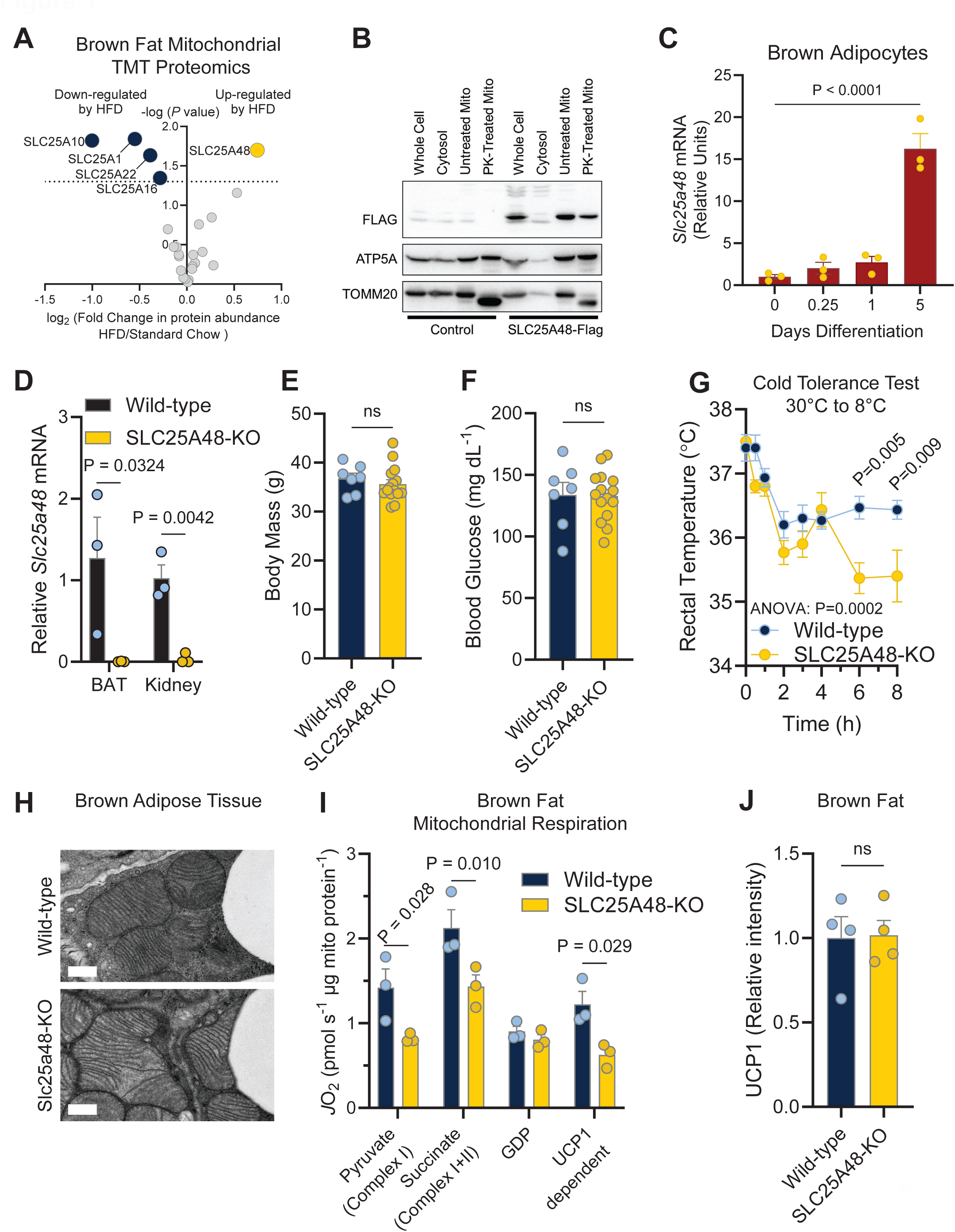
SLC25A48 is required for optimal brown adipose tissue thermogenesis. **A.** SLC25A family protein abundance from brown adipose tissue (BAT) mitochondria of wild-type mice fed a standard diet or high fat diet for 8 weeks. Mitochondria were isolated using differential centrifugation and subjected to quantitative tandem mass tag proteomics. N = 5 per group. Statistic: unpaired t-test. Yellow represents statistically upregulated by high fat diet and blue represents statistically down-regulated by high fat diet feeding. **B.** Proteinase K (PK) assay in brown adipocyte mitochondria. Mitochondria from brown adipocytes expressing control (empty-vector) or SLC25A48-Flag cDNA were exposed to a timed Proteinase K degradation to selectively degrade proteins on the outer mitochondrial membrane (OMM) and probed for OMM marker TOMM20, inner mitochondrial maker (IMM) ATP5A, and SLC25A48 through Flag. In control and SLC25A48-Flag the OMM marker TOMM20 was selectively degraded in PK treated mitochondria, while the IMM marker (ATP5A) was unchanged. Consistent with localization to the IMM, SLC25A48-Flag protein was not changed with PK treatment. **C.** Expression of SLC25A48 during the course of wild-type brown adipocyte differentiation. Wild-type brown adipocytes were collected at 0.25, 1, and 5 days after the addition of induction media (850 nM insulin, 1 nM T3, 125 nM indomethacin, 2 μg/mL dexamethasone, 0.5 mM isobutylmethylxanthine). After 2 days, cells were switched to maintenance media (850 nM insulin and1 nM T3) and were maintained until full differentiation at day 5. N = 3 per time point. Statistic: one-way ANOVA with Tukey’s multiple comparisons test. Bars represent mean and error shown as s.e.m. **D.** Confirmation of whole-body SLC25A48-knockout (KO) mouse in brown adipose tissue (BAT) and kidney. Germline knockout of SLC25A48 were generated through deletion of exon 4. Mice with heterozygous knockout of SLC25A48 were crossed to generate wild-type littermate controls and SLC25A48 homozygous knockout mice. Values relative to wild-type control. N = 3 per group. Statistic: unpaired t-test. Bars represent mean and error shown as s.e.m. **E.** Body mass of littermate control wild-type and SLC25A48-KO male mice at 6 months of age. N = 7 wild-type, 15 SLC25A48-KO. Statistic: unpaired t-test. Bars represent mean and error shown as s.e.m. **F.** Fasting blood glucose in standard diet fed adult male control (wild-type) and SLC25A48-KO mice. Mice were fasted for 4 hours prior to measurement of glucose via handheld glucometer. N = 7 wild-type, 15 SLC25A48-KO. Statistic: unpaired t-test with individual values shown. Bars represent mean and error shown as s.e.m. **G.** Cold tolerance test of standard diet fed male control (wild-type) and SLC25A48-KO mice. Mice were exposed to thermoneutral (30°C) for 10 days before being placed in cold 8°C chamber. N = 3 per group. Statistic: two-way ANOVA with Šídák’s multiple comparisons test. Bars represent mean and error shown as s.e.m. **H.** Brown adipose tissue mitochondrial structure of 12 week old control (wild-type) and SLC25A48-KO mice fed standard diet. Representative image. Scale bar = 0.5μm **I.** Brown adipose tissue isolated mitochondrial bioenergetics. Mitochondria were isolated from brown adipose tissue of control (wild-type) and SLC25A48-KO mice and stimulated in series with pyruvate (complex I mediated respiration), succinate (complex I+II mediated respiration), and guanosine diphosphate (GDP; inhibits UCP1). UCP1-dependent respiration was determined by subtracting GDP-respiration from complex I+II respiration (succinate). N = 3 per group. Statistic: two-way ANOVA with Šídák’s multiple comparisons test. Bars represent mean and error shown as s.e.m., individual values presented. **J.** Relative quantified protein abundance of UCP1 in brown fat from control and SLC25A48-KO mice. N = 4 per group. Statistic: unpaired t-test. Bars represent mean and error shown as s.e.m., individual values presented.

To better understand the role of SLC25A48 *in vivo,* we acquired mice with a germline knockout of SLC25A48 from the Knockout Mouse Phenotyping Program (KOMP). The knockout of SLC25A48 targeted exon 4, which is predicted to encode two of the six transmembrane domains (**Supplemental Fig. 1C**). In order to generate littermate controls, mice with heterozygous knockout of *Slc25a48* were crossed to generate littermate controls and mice with homozygous deletion of *Slc25a48* (SLC25A48-KO) (**Figure 1D, Supplemental Fig. 1D**). Male mice with a knockout of *Slc25a48* develop normally and did not have differences in body mass or fasting blood glucose compared to littermate controls (**Figure 1E, 1F**). Similarly, female mice with knockout of *Slc25a48* develop normally and do not have differences in body mass compared to littermate wild-type controls (**Supplemental Fig. 1E**). To determine if SLC25A48 has a role in brown adipose tissue function, we challenged mice with a cold exposure at 8°C. SLC25A48-KO mice displayed significantly impaired cold tolerance compared to littermate controls (**Figure 1G**). Although the gross BAT morphology appeared indistinguishable between the genotypes, electron microscopy analyses found that the mitochondrial ultrastructure of SLC25A48-KO BAT contained less dense cristae than control BAT (**Figure 1H**).

Impaired cold tolerance was due to a defect in BAT thermogenesis because mitochondrial respiration of isolated BAT via Complex I and II was significantly attenuated in both male and female SLC25A48-KO mice relative to littermate controls (**Figure 1I, Supplemental Fig. 1F**). In the presence of guanosine diphosphate (GDP), we did not observe differences in respiration rate, suggesting that UCP1-mediated thermogenesis was attenuated in the BAT of SLC25A48-KO mice. It is important to note, however, that the impairment in BAT thermogenesis was not due to down-regulation of UCP1 protein *per se* because UCP1 protein abundance was not different between the genotypes (**Figure 1J, Supplemental Fig. 1G**). Similarly, we did not find any difference in the protein levels of mitochondrial OXPHOS proteins between the two group (**Supplemental Fig. 1H**). Rather, genetic loss of SLC25A48 resulted in impaired Complex I and II activities thereby leading to reduced mitochondrial respiration and thermogenesis in BAT.

### Cell-autonomous regulation of choline and purine nucleotide metabolites by SLC25A48

We next aimed to address whether a defect in BAT thermogenesis of SLC25A48-KO mice arises from the cell-autonomous function of brown adipocytes. To this end, we established stable brown adipocyte cells from the stromal vascular fraction of SLC25A48-KO mouse brown adipose tissue (**Supplemental Fig. 2A**). Subsequently, we re-introduced a codon-optimized human SLC25A48-Flag cDNA to SLC25A48-KO immortalized cells to establish SLC25A48-rescued cells (SLC25A48-KO^SLC25A48-Flag^) (**Figure 2A**, **Supplemental Fig. 2B**). With these cells, we were able to exclude possible differences in cellular composition between control vs. SLC25A48-KO mice, and critically examine the cell-autonomous function of SLC25A48 using the identical origin of cells. There was no difference in brown adipogenesis between SLC25A48-KO cells and SLC25A48-rescued cells. Nonetheless, we found that norepinephrine-stimulated cellular respiration, both in total and oligomycin-independent respiration, was significantly lower in SLC25A48-KO cells than in SLC25A48-rescued cells (**Figure 2B**). Similar to brown adipose tissue, SLC25A48 loss did not alter mitochondrial protein abundance (**Supplemental Fig. 2C**); however, FCCP-induced maximum respiratory capacity was significantly lower in SLC25A48-KO cells relative to SLC25A48-rescured cells. Extracellular acidification rate was not different between the two groups, suggesting that impairments in cellular respiration are not due to changes in fuel preference (**Supplemental Fig. 2D**). Together, these results suggest that the requirement of SLC25A48 for mitochondrial respiratory activity is cell-autonomous.

**Figure 2.**
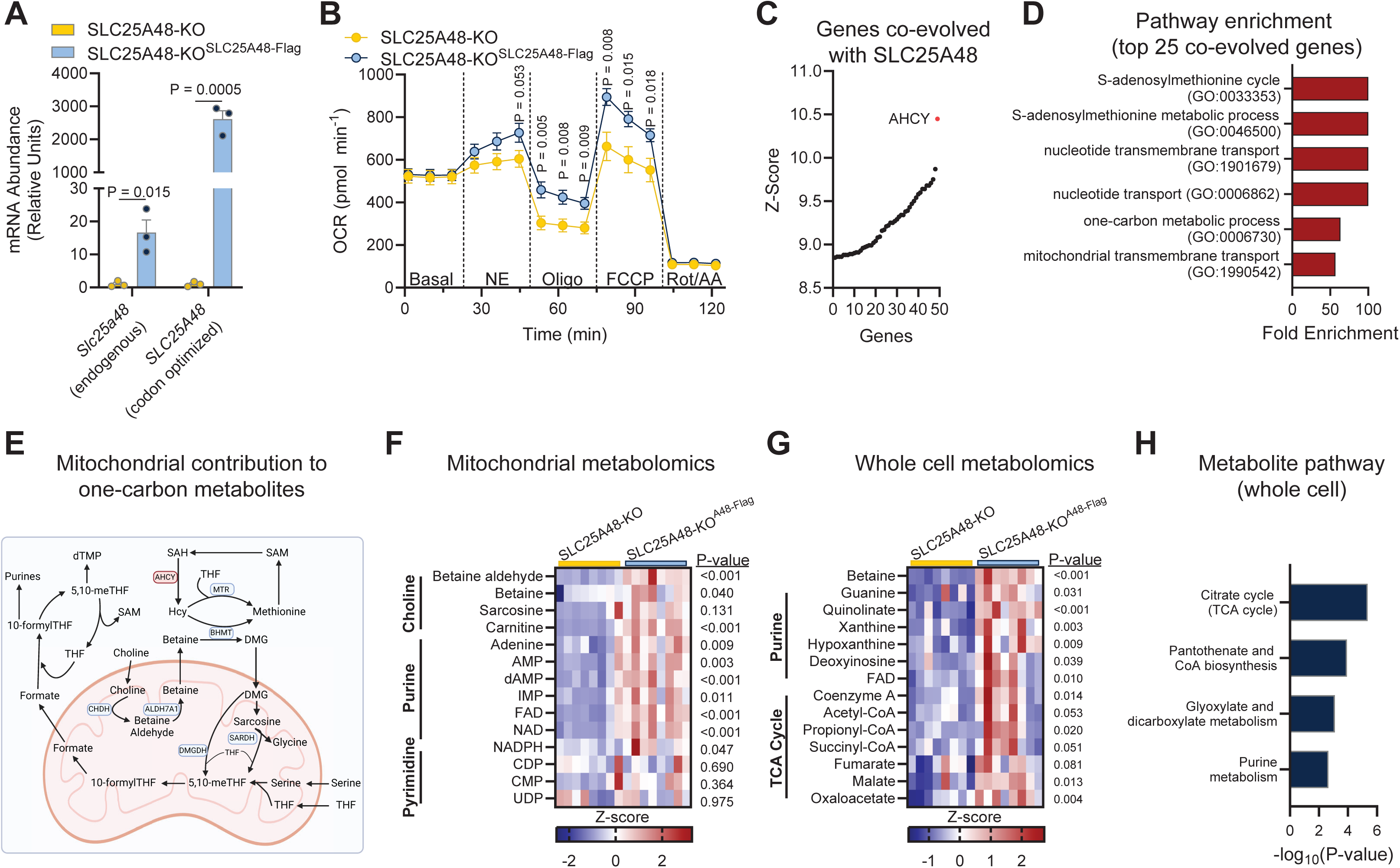
Cell-autonomous regulation of choline and purine nucleotide metabolites by SLC25A48. **A.** Expression of SLC25A48 (endogenous) and SLC25A48-Flag (Human cDNA) in SLC25A48-KO brown adipocytes and SLC25A48-KO brown adipocytes expression SLC25A48-Flag cDNA (SLC25A48-KO^SLC25A48-Flag^). Values relative to SLC25A48-KO. N=3 per group. Statistic: unpaired t-test. Bars represent mean and error shown as s.e.m., individual values presented. **B.** Cellular respiration of SLC25A48-KO and rescue (SLC25A48^SLC25A48-Flag^) brown adipocytes in response to norepinephrine (NE), oligomycin (oligo), carbonyl cyanide-p-trifluoromethoxyphenylhydrazone (FCCP), and rotenone and antimycin A (Rot/AA). N = 10 per group. Statistic is unpaired t-test. Circles represent mean and error shown as s.e.m. **C.** Top 50 genes co-evolved with human SLC25A48 across species ranked by z-score. Phylogene analysis of genes with similar phylogenetic profiles to SLC25A48. AHCY, adenosylhomocysteinase. **D.** Gene ontology pathway enrichment analysis for the top 25 genes to be co-evolved with SLC25A48 (Figure 2D). **E.** Schematic of mitochondrial contribution to one-carbon metabolites. Choline requires transport into the mitochondria to be metabolized by choline dehydrogenase (CHDH) and aldehyde dehydrogenase 7 family member A1 (ALDH7A1) to form one-carbon methyl donor betaine (trimethyl-glycine). Methyl-groups from betaine can be transferred to the methionine cycle or the folate cycle. Downstream metabolites that can be regulated by these pathways are purine nucleotides. Tetrahydratefolate (THF), 5,10-methylene-THF (5,10-meTHF), s-adenosylmethionine (SAM), s-adenosylhomocysteine (SAH). **F.** Mitochondrial metabolomics from brown adipocytes. Pathway enrichment of SLC25A48 suggests a role in methyl-donor into one carbon cycle and nucleotide homeostasis (Figure 1E). Metabolites that require import into mitochondria to contribute to one-carbon metabolism pathways include choline. Choline related metabolites are less abundant in SLC25A48-KO mitochondria. Methyl-groups from choline are an important precursor for the synthesis of purine nucleotides, which are less abundant in SLC25A48-KO mitochondria. Pyrimidine nucleotides were not different between SLC25A48-KO and SLC25A48-KO^SLC25A48-Flag^ mitochondria. FAD, flavin adenine dinucleotide; NADPH, nicotinamide adenine dinucleotide phosphate; NAD, nicotinamide adenine dinucleotide; dAMP, deoxyadenosine monophosphate; AMP, adenosine monophosphate; IMP, inosine monophosphate; CDP, cytidine diphosphate; CMP, cytidine monophosphate; UDP, uridine diphosphate. N = 8 per group. Statistic: unpaired t-test. Data represented as z-score. **G.** Whole cell metabolomics of SLC25A48-KO and SLC25A48-KO^SLC25A48-Flag^ brown adipocytes. Betaine was the most significantly less-abundant metabolite in SLC25A48-KO brown adipocytes compared to SLC25A48-KO^SLC25A48-Flag^ brown adipocytes. N = 8 per group. Statistic: unpaired t-test. Data represented as z-score. **H.** Pathway analysis of metabolites less abundant in whole cell of SLC25A48-KO brown adipocytes.

How does SLC25A48 regulate mitochondrial respiratory capacity? To address this question, we employed phylogene analysis (Sadreyev et al., 2015) to examine genes that co-evolved with SLC25A48 across species (**Figure 2C**). This analysis revealed adenosylhomocysteinase (AHCY) as the top co-evolved gene. AHCY is a cytosolic protein that is involved in one-carbon methionine cycle and produces a reversible hydrolysis reaction producing homocysteine and adenosine from s-adenosylhomocysteine (SAH). To analyze the co-evolved genes in more detail, we performed a gene ontology analysis of the top 25 co-evolved genes (**Figure 2D**). The highly enriched pathways involved the one-carbon cycle pathway that is well-known to play an important role in the formation of nucleotides (Petrova et al., 2023). Of note, one-carbon units in mammalian cells come from choline, serine, and glycine, all of which are imported into the mitochondria for the synthesis of purine nucleotides (**Figure 2E**) (Ducker and Rabinowitz, 2017). For instance, serine is previously shown to be imported into the mitochondrial matrix by sideroflexin 1 (SFXN1) (Kory et al., 2018), while the import of glycine is mediated by SLC25A38 (Fernandez-Murray et al., 2016). Similarly, choline must be imported into the mitochondrial matrix for the catabolic process because choline dehydrogenase (CHDH) and aldehyde dehydrogenase 7 family member A1 (ALDH7A1) are localized in the mitochondrial matrix to form the methyl donor metabolite betaine. However, the underlying mechanisms of choline import into the mitochondria and betaine export into the cytosolic compartment remain poorly characterized.

To determine if SLC25A48 controls the one-carbon metabolic pathway, we performed LC-MS metabolomics in whole-cells and isolated mitochondria from SLC25A48-KO and SLC25A48-rescued cells. Here, we employed the MITO-Tag expression system, which enabled us to perform rapid mitochondrial isolation and metabolite characterization by LC-MS (Chen et al., 2016). In isolated mitochondria, we found that choline degradation metabolites, including betaine aldehyde, betaine, and sarcosine, were significantly less in SLC25A48-KO cells relative to SLC25A48-rescued cells (**Figure 2F**). Moreover, SLC25A48-KO mitochondria contained significantly less amounts of purine nucleotides, such as adenine, adenosine monophosphate (AMP), dAMP, inosine monophosphate (IMP), and flavin adenine dinucleotide (FAD) (**Figure 2F**). Likewise, mitochondrial contents of nicotinamide adenine dinucleotide (NAD) and nicotinamide adenine dinucleotide phosphate (NADPH) were significantly lower in SLC25A48-KO cells compared to SLC25A48-rescued cells. The change in nucleotide levels was selective to purines as we did not find differences in pyrimidines, such as cytidine diphosphate (CDP), cytidine monophosphate (CMP), and uridine diphosphate (UDP). Importantly, the changes in mitochondrial metabolites immensely impact whole-cell metabolite profile: betaine was the most reduced metabolite in SLC25A48-KO cells. Furthermore, several other purine metabolites, such as guanine, quinolate, xanthine, hypoxanthine, deoxyinosine, and FAD, were significantly lower in SLC25A48-KO cells than SLC25A48-rescued cells (**Figure 2F**). In addition to purine metabolism, the pathway analysis of SLC25A48-regulated metabolites suggests that TCA cycle, CoA biosynthesis, and glyoxylate and dicarboxylate metabolism were down-regulated in SLC25A48-KO cells (**Figure 2H**). This is consistent with and may explain reduced mitochondrial Complex I and II respiration of SLC25A48-KO brown adipocytes (**Figure 2B**). Together, the whole-cell and mitochondrial metabolomics studies strongly implicate SLC25A48 in the regulation of choline metabolism.

### SLC25A48 is required for mitochondrial choline import and catabolism

The above data, *i.e.,* choline-derived metabolites are attenuated in SLC25A48-KO mitochondria, suggest that SLC25A48 is involved in mitochondrial choline import. To test the hypothesis, we utilized heavy-labeling of choline tracing coupled with whole-cell and mitochondrial metabolomics (**Figure 3A**). Here, we incubated SLC25A48-KO and SLC25A48-rescued brown adipocytes with D9-choline for 24 hours, collected cells and split into a whole-cell fraction and a mitochondrial-enriched fraction using differential centrifugation. In the whole-cell, both labeled (D9, red) and unlabeled (D0, blue) were significantly less abundant in SLC25A48-KO brown adipocytes compared to SLC25A48-rescued cells (**Figure 3B**). Similarly, labeled fractions of choline-derived metabolites betaine aldehyde (**Figure 3C**) and betaine (**Figure 3D**) were also significantly lower in whole-cell fraction of SLC25A48-KO cells than that of SLC25A48-rescued cells. In the mitochondrial fraction, choline import into the mitochondria was significantly less in SLC25A48-KO cells than SLC25A48-rescued cells (**Figure 3E**). Notably, D9-labeled betaine aldehyde in the mitochondria was nearly undetected in SLC25A48-KO cells, whereas its level was substantially elevated in the mitochondria of SLC25A48-rescued cells (**Figure 3F**). Furthermore, unlabeled-betaine aldehyde was not detected in both whole-cell (**Figure 3C**) and the mitochondrial fraction (**Figure 3F**), suggesting that SLC25A48 is required for the synthesis of choline-derived betaine aldehyde in the mitochondria. It is also worth noting that D9-labeled betaine was elevated in whole-cell (**Figure 3D**) of SLC25A48-rescued cells relative to SLC25A48-KO cells, even though there was no difference in the mitochondrial compartment (**Figure 3G**). The data suggest that SLC25A48 may also be involved in betaine export from the mitochondria to the cytosolic compartment. Independent from our work, we note here that two recent pre-prints deposited in medRxiv and bioRxiv state that SLC25A48 is involved in the mitochondrial choline transport (Khan et al., 2023; Patil et al., 2023). Together, these results further support the conclusion that SLC25A48 controls mitochondrial choline transport and catabolism.

**Figure 3.**
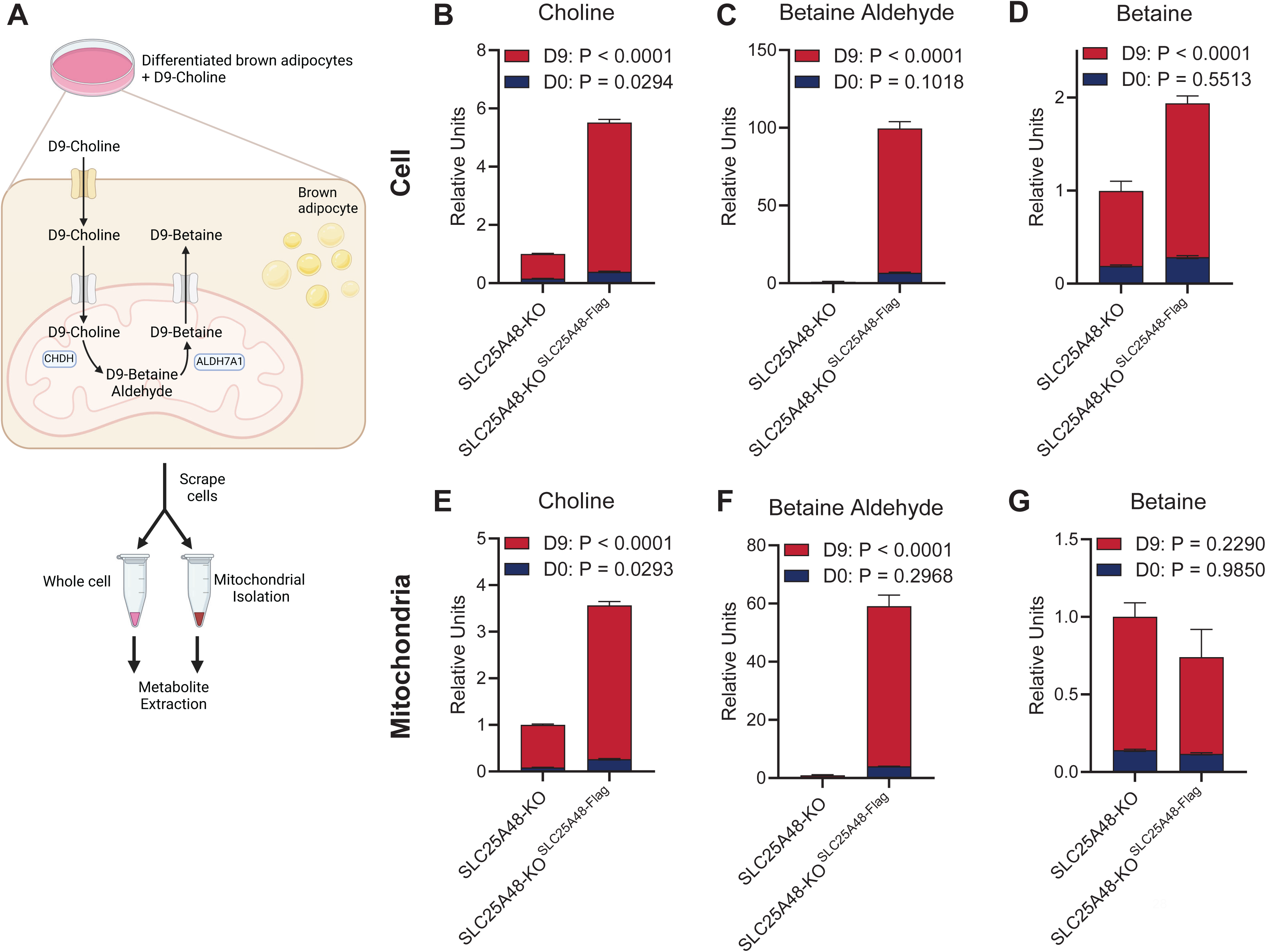
SLC25A48 is required for mitochondrial choline import and catabolism. **A.** Schematic of mitochondrial choline transport tracing assay. In the mitochondria, choline is converted to betaine aldehyde by choline dehydrogenase (CHDH) which is further processed to betaine by aldehyde dehydrogenase 7 family member A1 (ALDH7A1). Differentiated brown adipocytes were treated with 1 mM D9-choline for 24 hours. After 24 hours, cells were washed, scraped, and split into whole cell fraction and a fraction designated for mitochondrial isolation through differential centrifugation. Metabolites were extracted from whole cell and mitochondrial fractions and subjected to mass spectrometry analysis of choline and its metabolites. **B.** Whole cell labeled (D9, red) and unlabeled (D0, blue) choline in SLC25A48-KO and SLC25A48-KO^SLC25A48-Flag^ brown adipocytes. Statistic: two-way ANOVA with Šídák’s multiple comparisons test. Values relative to total choline (D0 + D9) of SLC25A48-KO. Bars represent mean and error shown as s.e.m. **C.** Whole cell labeled (D9, red) and unlabeled (D0, blue) betaine aldehyde in SLC25A48-KO and Slc25a48-KO^SLC25A48-Flag^ brown adipocytes. Statistic: two-way ANOVA with Šídák’s multiple comparisons test. Values relative to total betaine aldehyde (D0 + D9) of SLC25A48-KO. Bars represent mean and error shown as s.e.m. **D.** Whole cell labeled (D9, red) and unlabeled (D0, blue) betaine in SLC25A48-KO and SLC25A48-KO^SLC25A48-Flag^ brown adipocytes. Statistic: two-way ANOVA with Šídák’s multiple comparisons test. Values relative to total choline (D0 + D9) of SLC25A48-KO. Bars represent mean and error shown as s.e.m. **E.** Mitochondrial labeled (D9, red) and unlabeled (D0, blue) choline in SLC25A48-KO and SLC25A48-KO^SLC25A48-Flag^ brown adipocytes. Statistic: two-way ANOVA with Šídák’s multiple comparisons test. Values relative to total choline (D0 + D9) of SLC25A48-KO. Bars represent mean and error shown as s.e.m. **F.** Mitochondrial labeled (D9, red) and unlabeled (D0, blue) betaine aldehyde in SLC25A48-KO and SLC25A48-KO^SLC25A48-Flag^ brown adipocytes. Statistic: two-way ANOVA with Šídák’s multiple comparisons test. Values relative to total betaine aldehyde (D0 + D9) of SLC25A48-KO. Bars represent mean and error shown as s.e.m. **G.** Mitochondrial labeled (D9, red) and unlabeled (D0, blue) betaine in SLC25A48-KO and SLC25A48-KO^SLC25A48-Flag^ brown adipocytes. Statistic: two-way ANOVA with Šídák’s multiple comparisons test. Values relative to total choline (D0 + D9) of SLC25A48-KO. Bars represent mean and error shown as s.e.m.

In contrast, the following experiment suggests that SLC25A48 is not involved in mitochondrial FAD import even though SLC25A48-KO mitochondria had lower levels of FAD compared to SLC25A48-rescued cells (see Figure 2F). First, we treated SLC25A48-KO and rescued cells with ^13^C_4_,^15^N_2_-Riboflavin (M+6) for 4 hours before collecting metabolomics in whole-cell and mitochondrial fractions (**Supplemental Fig. 3A**). In whole-cell fraction, we found a trend of lower unlabeled (M+0) FAD levels in SLC25A48-KO cells, whereas the labeled fraction (M+9) was unchanged (**Supplemental Fig. 3B**). In the mitochondrial fraction, unlabeled FAD was also significantly less, while labeled-FAD levels were not different between the two groups (**Supplemental Fig. 3C**). This suggests the lower abundance of mitochondrial FAD in the SLC25A48-KO cells is likely due to reduced synthesis of purine nucleotides, rather than changes in the flux of riboflavin to FAD or the mitochondrial import of FAD.

### Mitochondrial choline catabolism via SLC25A48 regulates cell proliferation and survival

SLC25A48 mediates choline uptake into the mitochondria, allowing for choline catabolism in the mitochondria and the synthesis of the downstream metabolites, such as betaine and purine nucleotides. This is in alignment with the fact that choline metabolic byproducts betaine, dimethylglycine, and sarcosine are one-carbon units that feed into methionine and folate cycles (Ducker and Rabinowitz, 2017). One carbon metabolism and the purine nucleotide synthesis play a key role in cell cycle and the rate of proliferation; hence, we next aimed to determine the extent to which mitochondrial choline catabolism by SLC25A48 could alter cell proliferation. To this end, we developed stable HEK 293T cell lines using CRISPR/Cas9-mediated gene knockout of control and SLC25A48-KO. Subsequently, we re-expressed a codon-optimized SLC25A48-Flag cDNA in SLC25A48-KO cells at a level equivalent to endogenous SLC25A48 expression (**Supplemental Fig. 4A**). Consistent with brown adipocytes, we found that SLC25A48 is required for the synthesis of mitochondrial choline-derived metabolites: CRISPR/Cas9-mediated SLC25A48 loss reduced the levels of choline-derived metabolites, whereas re-expression of SLC25A48 in SLC25A48-KO cells restored the levels of betaine, betaine aldehyde, GDP, and NAD (**Figure 4A**, **Supplemental Fig. 4B**). Importantly, impaired synthesis of choline-derived metabolites in SLC25A48-KO cells was accompanied by reduced cell growth, while re-expression of SLC25A48 in SLC25A48-KO cells was sufficient to completely restore cell proliferation equivalent to wild-type cells (**Figure 4B**).

**Figure 4.**
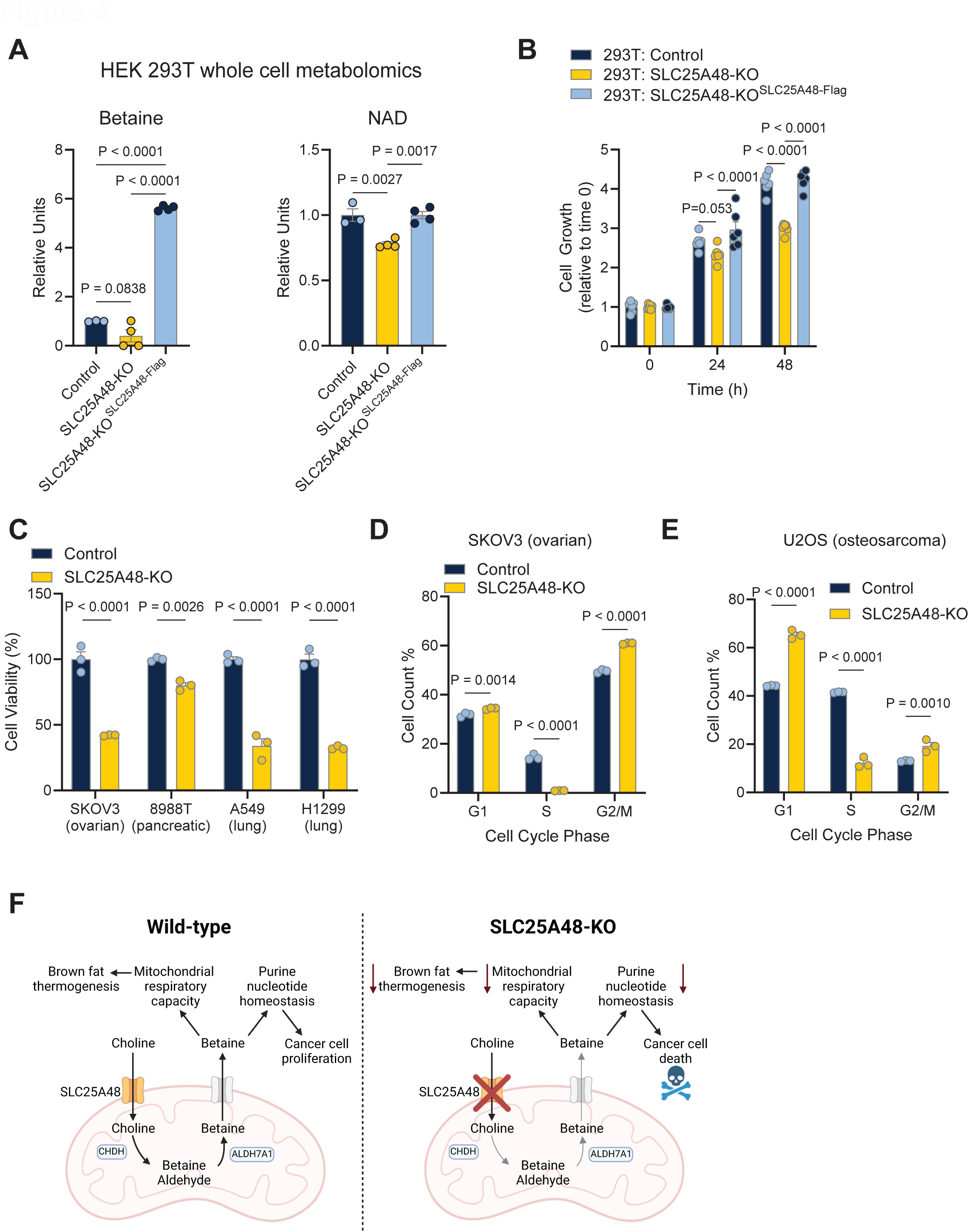
Mitochondrial choline catabolism via SLC25A48 regulates cell proliferation and survival. **A.** Whole cell metabolite abundance of betaine and nicotinamide adenine dinucleotide (NAD) of control, SLC25A48-KO, and SLC25A48-KO^SLC25A48-Flag^ HEK 293T cells. N = 3 per goup. Statistic: one-way ANOVA with Tukey’s multiple comparisons test. Bars represent mean and error shown as s.e.m., individual values presented. **B.** Cell growth in HEK-293T cells is impaired by loss of SLC25A48. Control, SLC25A48-KO, and SLC25A48-KO^SLC25A48-Flag^ HEK 293T cells cell growth over 48 hours, relative to time 0 h. N = 6 per group. Statistic: two-way ANOVA with Tukey’s multiple comparisons test. Bars represent mean with error shown as s.e.m., individual values presented. **C.** Cell viability of ovarian, pancreatic, and lung cancer cell lines in response to knockout of SLC25A48. Cells were transfected with control or SLC25A48-KO plasmid and cell viability was measured via MTS assay 24 hours post transfection. N= 3 per group. Statistic: two-way ANOVA with Šídák’s multiple comparisons test. Bars represent mean with error shown as s.e.m., individual values presented. **D.** Cell cycle analysis of SKOV3 ovarian cancer cells 16 hours after transfection with control or SLC25A48-KO plasmid. N= 3 per group. Statistic: two-way ANOVA with Šídák’s multiple comparisons test. Bars represent mean with error shown as s.e.m., individual values presented. **E.** Cell cycle analysis of U2OS osteosarcoma cancer cells 16 hours after transfection with control or SLC25A48-KO plasmid. N= 3 per group. Statistic: two-way ANOVA with Šídák’s multiple comparisons test. Bars represent mean with error shown as s.e.m., individual values presented. **F.** Mitochondrial choline import regulates purine nucleotide pools through SLC25A48. Choline import into mitochondria is required for the conversion of choline into its downstream metabolites. The import of choline into the mitochondria regulates purine nucleotide pools, mitochondrial respiratory capacity, and thermogenesis in brown adipose tissue. In cancer cells, mitochondrial choline import is required for cell proliferation.

Given the critical role of *de novo* purine nucleotide synthesis in cancer (Mullen and Singh, 2023), we then asked if SLC25A48-mediated changes in cell proliferation extended to cancer cells. Here, we used four independent human cancer cells – ovarian cancer cells (SKOV3), pancreas adenocarcinoma (PA-TU-8988T), non-small cell lung cancer cell lines (A549 and H1299). In all cancer cells tested, genetic loss of SLC25A48 by the CRISPR/Cas9 system resulted in a significant reduction in cell viability (**Figure 4C, Supplemental Figure 4C**). Within 24 hours post induction of SLC25A48 knockout, we found over 50% of ovarian and lung cancer cells died, and 20% of pancreatic cancer cells were dead. Since loss of SLC25A48 leads to reduced pool sizes of mitochondrial choline-derived purine nucleotides, we next tested if SLC25A48-deletion impairs cell cycle progression. We found that ovarian cancer cells lacking SLC25A48 failed to initiate the G1-to-S phase transition (**Figure 4D**). Similarly, knockout of SLC25A48 in U2OS osteosarcoma cells impaired cell cycle in the transition from G1-to-S phase (**Figure 4E**). Together, these data suggest that SLC25A48-mediated choline import is required for cancer cell cycle progression, likely through mitochondria choline catabolism contributions to purine nucleotide synthesis.

## DISCUSSION

This present study suggests a model (**Figure 4F**) in which SLC25A48 mediates the import of choline into the mitochondrial matrix to allow for the conversion of choline to downstream metabolites, including a critical methyl-donor, betaine. Our data show that this SLC25A48-mediated pathway is necessary for mitochondrial respiratory capacity (Complex I and II activity) and purine nucleotide homeostasis. It is worth noting that the mitochondrial phenotype of SLC25A48-KO brown fat resembles that of mice lacking CHDH – the first catabolic enzyme of choline in the mitochondria, which exhibits poor mitochondrial cristae density, low ATP levels, and less mitochondrial membrane potential in the knockout sperm (Johnson et al., 2010). We initially identified SLC25A48 as a uniquely upregulated IMM-bound protein in BAT following high-fat diet feeding and also treatment with a committed differentiation cocktail. This regulation is intriguing as SLC25A48 may have implications for lipid metabolism. Choline has been well-appreciated as an important nutrient for the synthesis of phospholipids, such as phosphatidylcholine and sphingomyelin. A deficiency in choline metabolism is linked to hepatic steatosis and liver diseases in which the expression of phosphatidylethanolamine (PE) methyltransferase (PEMT) increases in the liver in response to obesity (Fu et al., 2011; Song et al., 2005). The role of SLC25A48 in the pathogenesis of fatty liver disease remains unknown; thus, future studies incorporating liver-specific models will determine the role of mitochondrial choline metabolism in the liver with the pathogenesis of fatty liver disease.

The present work provided important insights into mitochondrial choline catabolism because a choline transporter-like protein 1 (CLT1, *SLC44A1*), which was initially characterized as a plasma-membrane choline uptake (O’Regan et al., 2000; Yuan et al., 2004), was implicated as a mediator of mitochondrial choline transport (Michel and Bakovic, 2009). However, the data analysis in MitoCarta 3.0 (Rath et al., 2021) found no detection of SLC44A1 in the mitochondria. On the other hand, our tracing study showed the functional requirement of SLC25A48 for mitochondrial choline import and choline catabolism, including the synthesis of betaine in the mitochondrial compartment. Nevertheless, we are aware that labeled choline was still detected in the mitochondrial fraction of SLC25A48-KO cells, albeit to a substantially lesser extent. This suggests the possibility of redundant mitochondrial choline carriers. In fact, several metabolites have redundant carriers for the import into the mitochondrial matrix, including adenosine diphosphate (ADP) (*SLC25A4, SLC25A5, SLC25A6*), glutamate (*SLC25A12, SLC25A13, SLC25A18, SLC25A22*), pyrimidine nucleotides (*SLC25A33, SLC25A36*) and NAD (*SLC25A51, SLC25A52*), depending on tissue distribution of each carrier protein (Luongo et al., 2020; Palmieri, 2013). In this regard, SLC25A48 is highly enriched in the BAT, but not in the heart and soleus where redundant choline transporters may play a role. Another notable observation is that choline-derived betaine was accumulated in the mitochondria of SLC25A48-KO cells, despite labeled betaine levels being 2-fold more in the whole-cell of SLC25A48-rescue cells. The data suggests that SLC25A48 is also involved in betaine export from the mitochondria to the cytosolic compartment.

The critical role of SLC25A48 for the synthesis of methyl-donor betaine immediately implicated the possible role in cell proliferation. The *de novo* synthesis of purine nucleotides requires one-carbon donor, and diet-derived choline makes up nearly 60% of methyl-donors consumed (Niculescu and Zeisel, 2002). It is notable that choline is a high consumption metabolite across NCI-60 cell lines (Jain et al., 2012) even though the preferred source of one-carbon donor depends on cell type (Ducker and Rabinowitz, 2017) and serine are the preferred source of one-carbon units (Labuschagne et al., 2014). Regardless, we found that a defect in mitochondrial choline catabolism by the deletion of SLC25A48 and subsequent reduction in the purine nucleotide pool resulted in failed G1-to-S phase transition and cell death in several cancer cell lines. The present data suggest that limiting mitochondrial choline catabolism provides a unique approach to manipulating cell proliferation in some cancer cells. How the SLC25A48-mediated choline catabolic pathway is regulated in pathophysiology, including cancer and metabolic disorders, is an important area of future research.

## Acknowledgments

We thank Mark Li and Tadashi Yamamuro for their technical support, John Asara and the BIDMC mass spectrometry core facility for targeted metabolomic work. We thank Susan Hagen and Kyle Smith of the BIDMC electron microscopy core. This work was supported by grants from the National Institutes of Health (NIH) (DK125283, DK097441, DK126160) and Howard Hughes Medical Institute to SK. ARPV is supported by King Trust, Bank of America Private Bank, Co-Trustees.

## METHODS

### Animals

All animal experiments were performed in compliance with approved protocols by the Institutional Animal Care and Use Committee at Beth Israel Deaconess Medical Center. All mice were housed were housed under a 12 h light – 12 h dark cycle. Room temperature mice were housed at 23°C in ventilated cages with an ACH of 25. Thermoneutral housed mice, were placed in an incubator set to 30°C. Mice were fed a standard diet (Lab Diet 5008) and had free access to food and water. Mice were fasted 4 h prior to metabolic measurements of fasting blood glucose and mitochondrial respiration. Slc25a48-knockout mice were acquired from the Knockout Mouse Phenotyping Program (KOMP) at Jackson Laboratory (Strain# 051066-JAX) and backcrossed to wild-type mice (Jackson Laboratory, Strain# 000664). Slc25a48-heterozygous knockout mice were used for breeding littermate controls.

### Cell Culture

The stromal vascular fraction (SVF) from the brown adipose tissue of Slc25a48-KO mice were immortalized by expressing the SV40 large T antigen as previously described (Shinoda et al., 2015). The base media for adipocyte cell culture was DMEM (Gibco 11965092), containing 10% FBS and 1% penicillin/streptomycin (Gibco 15140). To induce differentiation to brown adipocytes, confluent preadipocytes were treated with an induction cocktail containing 0.5 mM isobutylmethylxanthine (Sigma I5879), 125 nM indomethacin (Sigma I7378), 2 μg/mL dexamethasone (Sigma D4902), 850 nM insulin (Sigma: I6634), and 1 nM T3 (Sigma T2877) for 2 days. After induction, cells were changed to a maintenance medium of base medium supplemented with 850 nM insulin and 1 nM T3. Cells were kept in a maintenance medium for 4-6 days to reach full differentiation.

### Plasmid and Virus production

Codon optimized SLC25A48 was cloned from Addgene plasmid #131995 and cloned into pMSCV-blasticidin (Addgene #75085) with Flag-tag on C-terminus. All constructs were confirmed by sequencing. pMXs-3XHA-EGFP-OMP25 was obtained for MITO-Tag experiments (Addgene #83356). CRISPR/Cas9 plasmids for control and SLC25A48-KO (Santa Cruz Biotechnology; sc-418922, sc-414730, sc-414730-HDR) were transfected with UltraCruz transfection reagent and plasmid transfection medium (Santa Cruz Biotechnology; sc-395739, sc-108062). For virus production HEK293T packaging cells were transfected with 10 μg of retroviral or lentiviral plasmid and the packaging constructs through calcium phosphate method. After 48 h, the viral supernatant was collected and filtered through a 0.45 μm filter. Immortalized preadipocytes were infected with viral supernatant supplemented with 10 μg ml^-1^ polybrene for 24 h. Stable cell lines were selected with puromycin (10 μg ml^-1^) or blasticidin (10 μg ml^-1^). Cells expressing MITO-Tag construct were sorted for equally low abundant EGFP-positive cells (Chen et al., 2017)

### Cell Proliferation and Viability

Cells were counted and equally plated in 96-well. Cell proliferation and viability were determined through MTS assay kit (Abcam, ab197010). Cell proliferation were normalized to time 0 (24 h post seeding). Cell viability was measured 24 h post induction of SLC25A48-KO.

### Cell Cycle

Hour 16 post-transfection control and SLC25A48-KO were incubated with 10 µM EdU (Click-iT Plus EdU Flow Cytometry Assay Kit, Thermo Fisher Scientific, USA) for 1 h in DMEM/F12 media. Subsequently, cells were fixed according to the manufacture’s instruction. FxCycle Violet Ready Flow Reagent (Thermo Fisher Scientific) was used to measure the total DNA content, following the manufacturer’s instructions. Cell population (%) was calculated as the frequency of parent. All the cells were analyzed using a FACS Aria II equipped with 100 mm nozzle diameter and CytoFLEX. FlowJo software (version 10.8.1) and CytExpert (version 2.4.0.28) were used for data analyses.

### Cold Tolerance

Mice were acclimated to 30°C for 10 days prior to being single housed at 8°C for 8 hours. The rectal temperature of mice were monitored hourly using TH-5 thermometer (Physitemp).

### Cellular Respiration

Oxygen consumption rate (OCR) and extracellular acidification rate in fully differentiated brown adipocytes was measured with the Seahorse XFe Extracellular Flux Analyzer (Agilent) in a 24-well plate. Experiment was run in seahorse assay media supplemented with 10 mM glucose, 2 mM glutamine, and 1 mM pyruvate. Cells were stimulated with 10 μM norepinephrine, 5 μM oligomycin, 5 μM carbonyl cyanide-p-trifluoromethoxyphenylhydrazone (FCCP), and 5 μM of both rotenone and antimycin A.

### MITO-Tag mitochondrial isolation

Following established protocols (Chen et al., 2017), mitochondria were immunoprecipitated from brown adipocytes. In 100 mm cell culture dishes of differentiated brown adipocytes that expressed MITO-Tag construct (3xHA-EGFP -OMP25), cells were washed twice with PBS. Cells were then scraped with KPBS (136 mM KCl, 10 mM KH2PO4, pH 7.25). Cells were pelleted by 1000 x g for 2 min at 4°C. Whole cell fraction was extracted directly into 80% MeOH, and stored at -80°C for Cells were resuspended in 1 mL of KPBS. Mitochondrial fraction was homogenized with Teflon pestle and glass tube. Homogenized sample was centrifuged at 1000 x g for 2 min at 4°C. Supernatant was then added to magnetic anti-HAbeads (Pierce, 88836) and nutate for 4 min at 4°C. Next, beads were collected by 1 min on magnetic rack. Sample was aspirated and beads were washed 3 times with KPBS. After beads were washed, 80% MeOH was added to extract metabolites and samples were stored at -80C overnight. Samples were then vortexed for 1 min and then centrifuged at 20000 x g for 10 min at 4°C. Supernatant was then transferred to clean tube and dried via speed vac. Dried metabolites were stored at -80°C for up to 1 week until resuspension in LC/MS grade H_2_O for LC/MS analysis.

### Mitochondrial uptake with heavy label metabolites

Differentiated brown adipocytes were incubated with 1 mM D9-choline (CIL, DLM-549-PK) for 24 h or 1 μM ^13^C_4_,^15^N_2_-Riboflavin (CIL, CNLM-8851-PK) for 4 h in standard differentiation maintenance media (4.5 g/L glucose DMEM, 10% FBS, 1% penicillin/streptomycin, 850 nM insulin, and 1 nM T3). After incubation cells were washed with PBS, scraped and split into whole cell fraction and mitochondrial isolation fraction. Mitochondrial fraction was homogenized in buffer containing 300 mM sucrose, 10 mM HEPES, 1 mM EGTA. Homogenized samples were centrifuged at 600 x g for 5 min at 4°C and supernatant was transferred to a new tube and pelleted at 10000 x g for 10 min at 4°C. Whole cell and mitochondrial fractions were normalized to protein content. Cell and mitochondrial pellets were stored at -80°C prior to metabolite extraction and LC/MS analysis.

### Metabolomics

Labeled metabolomics was carried out with HRHA orbitrap instruments (Exploris 240/ Orbitrap ID-X) coupled with Vanquish Horizon UHPLC system. Waters ACQUITY UPLC BEH Amide column at 25 °C (particle size, 1.7 μm; 100mm (length) × 2.1mm (i.d.)) was used for LC separation. Mobile phases A = 25mM NH_4_Ac and 25mM ammonium hydroxide in water, and B = 100% acetonitrile were used for LC separation. The linear gradient as follows: 95% B (0.0– 1 min), 95% B to 65% B (1–7.0 min), 65% B to 40% B (7.0–8.0 min), 40% B (8.0–9.0 min), 40% B to 95% B (9.0–9.1 min), then stayed at 95% B for 5.9 min. The flow rate was 0.4 mL/min. The sample injection volume was 2 μL for cell lysate and 5 μL for media. ESI source parameters were set as follows: spray voltage, 3500 V or −2800 V, in positive or negative modes, respectively; vaporizer temperature, 350 °C; sheath gas, 50 arb; aux gas, 10 arb; ion transfer tube temperature, 325 °C. The full scan was set as: orbitrap resolution, 60,000; maximum injection time, 100 ms; scan range, 70–1050 Da. The ddMS2 scan was set as: orbitrap resolution, 30,000; maximum injection time, 60 ms; top N setting, 6; isolation width, 1.0 m/z; HCD collision energy (%), 30; Dynamic exclusion mode was set as auto. The ^15^N labeling metabolomics was quantified by Compound Discoverer 3.3

### Mitochondrial respiration

Brown adipose tissue was homogenized in buffer containing 300 mM sucrose, 10 mM HEPES, 1 mM EGTA, 1 mg/mL fatty acid-free BSA at pH 7.2. BAT homogenate was centrifuged at 600 x g for 5 min at 4°C and supernatant was centrifuged at 10000 x g for 10 min at 4°C to pellet mitochondria. Pelleted mitochondria were resuspended in homogenization buffer without BSA and normalized for equal protein through BCA assay. Mitochondrial respiration was measured with Oroboros O2k oxygraphs. For respiration experiments mitochondria were in buffer Z (100 mM MES potassium salt, 30 mM KCl, 10 mM KH_2_PO_4_, 5 mM MgCl_2_, 1 mM EGTA, 0.5 mg/ml fatty-acid free BSA pH 7.4). BAT mitochondria from male mice were stimulated with 0.5 mM malate, 5 mM pyruvate, and 10 mM succinate. BAT mitochondria from female mice were stimulated with 0.5 malate, 5 mM pyruvate, 5 mM glutamate, and 10 mM succinate. UCP1 was inhibited in both male and female mice with 4 mM GDP.

### Mitochondrial membrane localization

Cells were trypsinized and pelleted at 1100 x g for 2 min. Cell pellets were washed with phosphate buffer solution (PBS) and re-pelleted at 1100 x g for 2 min. Cell pellets were then homogenized in solution containing 22 mM mannitol, 75 mM sucrose, 1 mM EGTA, 30 mM Tris-HCl pH 7.4. Whole cell homogenate was split for whole cell fraction. Mitochondrial fraction was spun at 600 x g for 5 min and supernatant was pelleted for mitochondria at 7000 x g for 10 min. Assay was performed in 150 mM KCl, 10 mM HEPES, 200 μM CaCl_2_ buffer at pH 7.2. To mitochondrial fraction 0.25 μg of proteinase K was added and samples were incubated on ice for 10 min. After 10 min, Laemmli sample buffer was added and samples were boiled at 95°C for 10 min.

### Proteomics

Isolated mitochondrial proteomes were reduced with 5 mM TCEP, alkylated with 10 mM Iodoacetamide, further reduced with 5 mM DTT, and precipitated using TCA (final concentration of 20%). The precipitated samples were washed three times with ice-cold acetone. Proteins were solubilized in a digestion buffer (1M urea, 200 mM EPPS pH 8.5) and digested with Lys-C overnight. The samples were further digested with trypsin for 6 hours at 37°C for six hours. The digested samples were labeled with TMTPro reagents (Thermo Fisher Scientific). Following incubation at room temperature for 2 hours, the reactions were quenched with hydroxylamine to a final concentration of 0.5% (v/v). Following TMT labeling, the samples were combined, and the pooled sample was de-salted using a Sep-pak. To perform mitochondrial TMT proteomics, labeled peptides were fractionated using Pierce High pH Reversed-Phase Peptide Fractionation Kit (Thermo Scientific). A total of 6 fractions were collected. Samples were subsequently acidified with 1% formic acid and vacuum centrifuged to near dryness. Each consolidated fraction was desalted by StageTip, and reconstituted in 5% acetonitrile, 5% formic acid for LC-MS/MS analysis. Data were collected on an Orbitrap Fusion Lumos Tribird mass spectrometer (Thermo Fisher Scientific) equipped with a Thermo Easy-nLC 1000 for online sample handling and peptide separations. The 100 µm capillary column was packed in-house with 35 cm of Accucore 150 resin (2.6 μm, 150Å; ThermoFisher Scientific). The peptides were separated using a 180 min linear gradient from 5% to 32% buffer B (90% ACN + 0.1% formic acid) equilibrated with buffer A (5% ACN + 0.1% formic acid) at a flow rate of 550 nL/min across the column. Data was collected using an SPS-MS3 method. The scan sequence for the Fusion Lumos Orbitrap began with an MS1 spectrum collected in the Orbirap (resolution - 120,000; scan range - 350 – 1,500 m/z; AGC target – 1,000,000; normalized AGC target – 250%; maximum ion injection time 50 ms; dynamic exclusion - 180 seconds). MS2 spectra were collected in the ion trap following collision-induced dissociation (AGC target – 15,000; normalized AGC target - 150%; NCE (normalized collision energy) – 35; isolation window - 0.5 Th; maximum injection time - 50ms). MS3 scans were collected in the Orbitrap following higher-energy collision dissociation (resolution – 50,000; AGC target – 100,000; normalized AGC target – 200%; collision energy – 55%; MS2 isolation window – 2; number of notches – 10; MS3 isolation window – 1.2; maximum ion injection time – 200 ms. Proteomic Data Analyses: Database searching included all entries from the mouse UniProt Database (downloaded in May 2021). The database was concatenated with one composed of all protein sequences for that database in the reversed order (Elias and Gygi, 2007). Raw files were converted to mzXML, and monoisotopic peaks were re-assigned using Monocle (Rad et al., 2021). Searches were performed with Comet (Eng et al., 2013) using a 50-ppm precursor ion tolerance and fragment bin tolerance of 0.02. TMTpro labels on lysine residues and peptide N-termini +304.207 Da), as well as carbamidomethylation of cysteine residues (+57.021 Da) were set as static modifications, while oxidation of methionine residues (+15.995 Da) was set as a variable modification. Peptide-spectrum matches (PSMs) were adjusted to a 1% false discovery rate (FDR) using a linear discriminant after which proteins were assembled further to a final protein-level FDR of 1% analysis (Huttlin et al., 2010). TMT reporter ion intensities were measured using a 0.003 Da window around the theoretical m/z for each reporter ion. Proteins were quantified by summing reporter ion counts across all matching PSMs. More specifically, reporter ion intensities were adjusted to correct for the isotopic impurities of the different TMTpro reagents according to manufacturer specifications. Peptides were filtered to exclude those with a summed signal-to-noise (SN) < 160 across all TMT channels and < 0.5 precursor isolation specificity. The signal-to-noise (S/N) measurements of peptides assigned to each protein were summed (for a given protein).

### Pathway enrichment

Gene ontology pathway enrichment (Ashburner et al., 2000; Gene Ontology et al., 2023) was used to analyze biological processes from the top 25 genes from human SLC25A48 phylogene co-evolved prediction analysis (Sadreyev et al., 2015). MetaboAnalyst 6.0 pathway analysis was employed to analyze whole cell metabolites less abundant in Slc25a48-KO compared to Slc25a48-rescued brown adipocytes (Xia et al., 2009).

### Immunoblotting

Brown adipose tissue and brown adipocyte homogenates were probed for proteins of mitochondrial oxidative phosphorylation (Abcam, ab110413) and UCP1 (Sigma-Aldrich, U6382).

### Quantitative RT-PCR (qPCR)

Total RNA was isolated from cells or tissue using Trizol (Invitrogen) according to manufacturer instructions. RNA was reverse transcribed using iScript cDNA synthesis kit (Biorad). PCR reactions were performed with Applied Biosystems QuantStudio 6 Flex using Sybrgreen (Biorad). Assays were performed in duplicate, and all results were normalized to 18S ribosomal RNA or 36B4. Values are relative to the mean of the control group. Primers used are listed in **Supplementary Table 1**.

### Data analyses and Statistics

Statistical analyses were performed using GraphPad Prism v10. All data are represented at mean ± s.e.m. Unpaired t-tests were used for two-group comparisons. One-way ANOVA with Tukey’s multiple comparison test or two-way ANOVA with Šídák’s multiple comparison test were used for experiments with multiple comparisons. The statistical test used and sample numbers for each experiment are specified in the figure legends. P < 0.05 was considered to be significant.

---

**Supplemental Figure 1.**
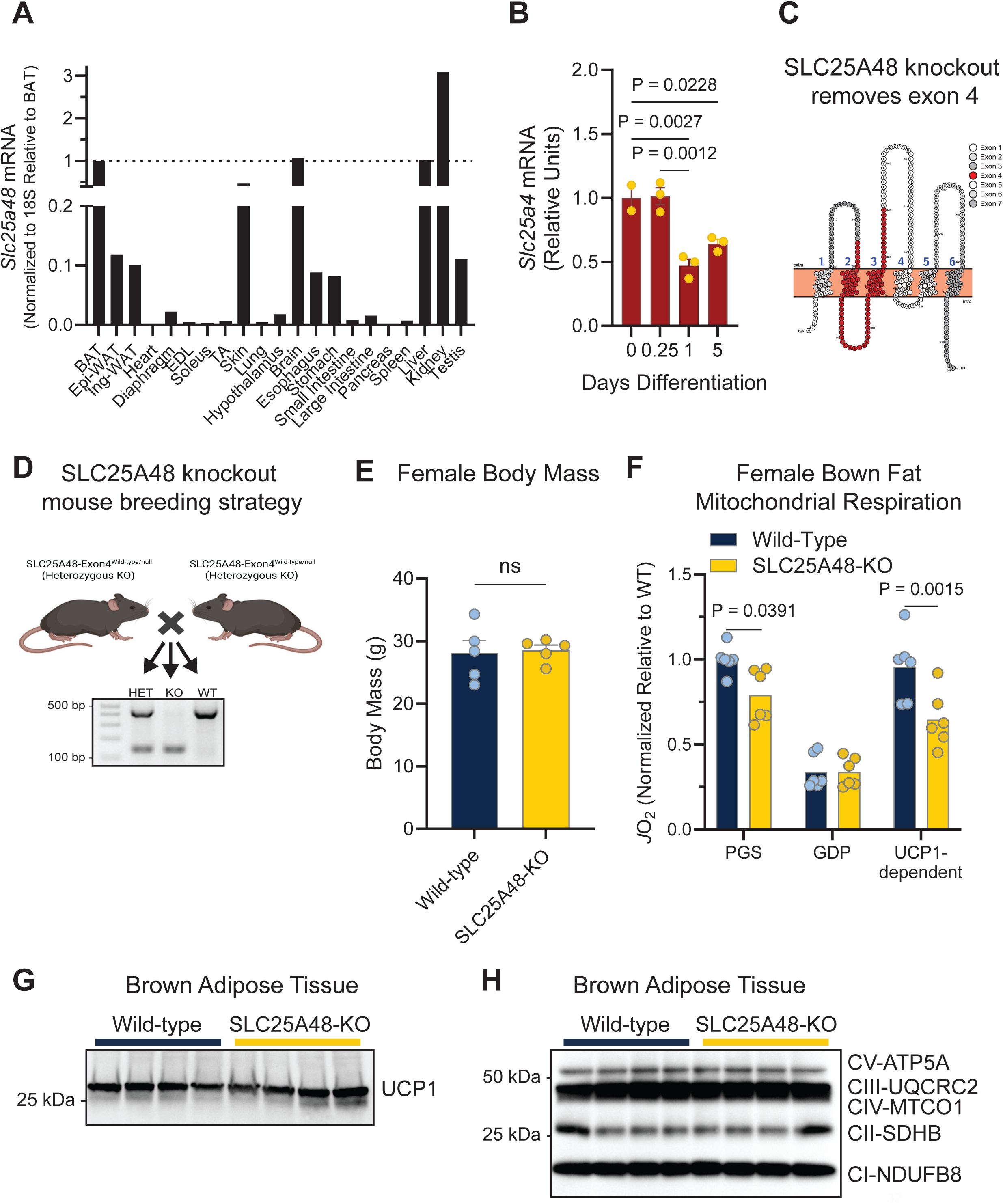
SLC25A48 is required for optimal brown adipose tissue thermogenesis. **A.** Expression of SLC25A48 in 6 week old wild-type mouse tissues. N = 1. BAT, brown adipose tissue; Epi-WAT, epididymal white adipose tissue; Ing-WAT, inguinal white adipose tissue; EDL, extensor digitorum longus; TA, tibialis anterior. **B.** Expression of SLC25A48 during the course of wild-type brown adipocyte differentiation. Wild-type brown adipocytes were collected at 0.25, 1, and 5 days after the addition of induction media (850 nM insulin, 1 nM T3, 125 nM indomethacin, 2 μg/mL dexamethasone, 0.5 mM isobutylmethylxanthine). After 2 days, cells were switched to maintenance media (850 nM insulin and1 nM T3) and were maintained until full differentiation at day 5. N = 3 per time point. Statistic: one-way ANOVA with Tukey’s multiple comparisons test. Bars represent mean and error shown as s.e.m. **C.** Amino acid sequence of mouse SLC25A48 highlighted by exon contribution. The strategy of SLC25A48-knockout mouse was to delete exon 4. Exon 4 is predicted to encode transmembrane domains 2 and 3 of SLC25A48. **D.** Breeding strategy for wild-type control and SLC25A48-KO mice. Mice with heterozygous deletion of SLC25A48 exon 4 were crossed to generate heterozygous knockout of SLC25A48 (HET), SLC25A48-KO (KO), or wild-type (WT) littermate controls. Wild-type and SLC25A48-KO mice were studied. **E.** Body mass of female wild-type control and SLC25A48-KO mice. N = 5 per group. Statistic: unpaired t-test. Bars represent mean and error shown as s.e.m. Individual values presented. **F.** Brown adipose tissue isolated mitochondrial bioenergetics in female mice. Mitochondria were isolated from brown adipose tissue of control (wild-type) and SLC25A48-KO mice and stimulated with pyruvate, glutamate, and succinate (PGS) to stimulate complex I and complex II respiration, followed by guanosine diphosphate (GDP) to inhibit UCP1. UCP1-dependent respiration was determined by subtracting GDP-respiration from complex I+II respiration (PGS). N = 6 per group. Statistic: two-way ANOVA with Šídák’s multiple comparisons test. Bars represent mean and error shown as s.e.m., individual values presented. **G.** Western blot for UCP1 in brown adipose tissue from male wild-type control and SLC25A48-KO mice. N = 4 per group. Western blot for mitochondrial complex proteins in brown adipose tissue mitochondria. Bands top to bottom complex IV (CV, ATP5A), complex III (CIII, UQCRC2) is overlapped with complex IV (MTCO1), complex II (SDHB), complex I (NDUFB8). N = 4 per group.

**Supplemental Figure 2.**
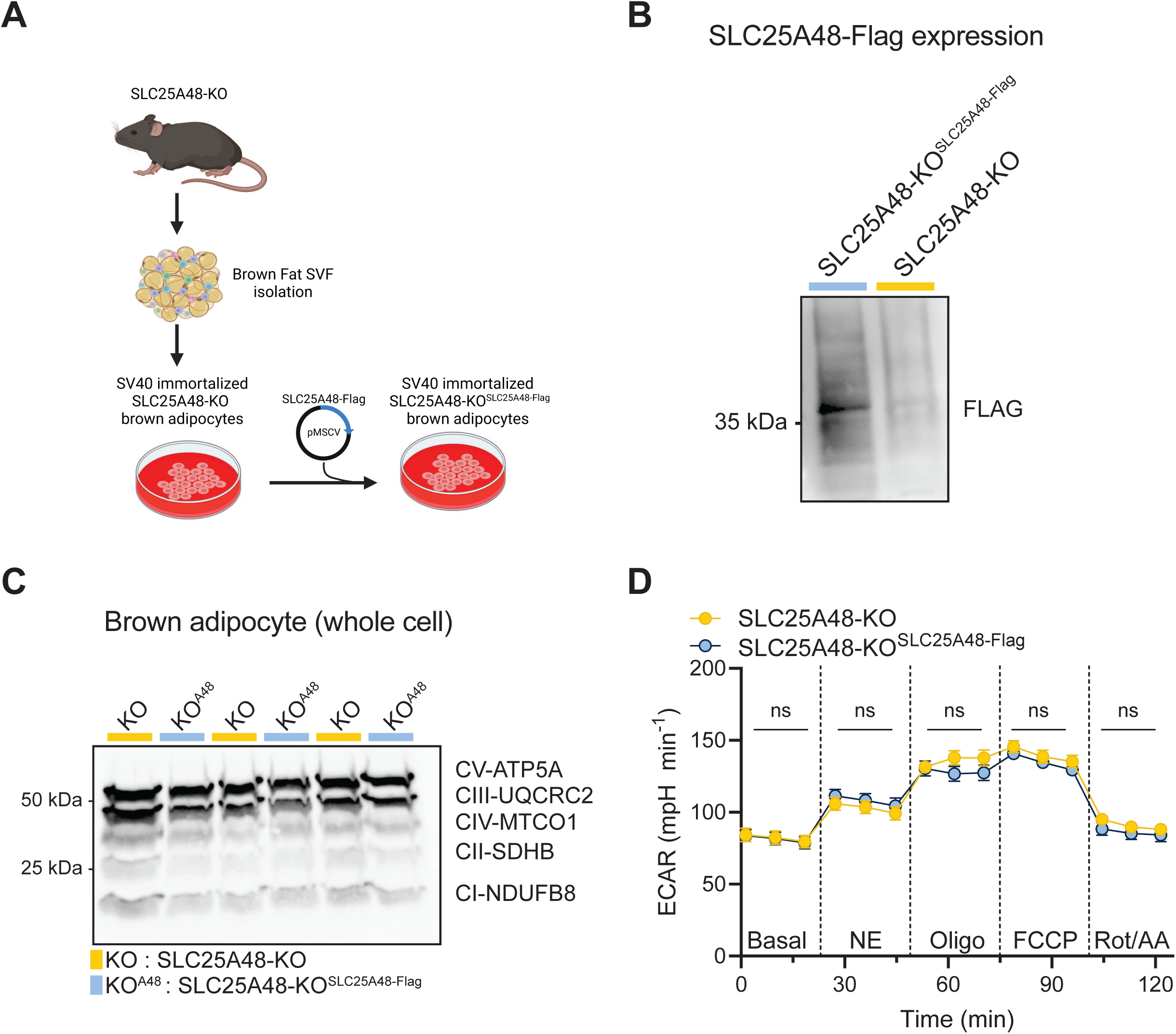
Cell-autonomous regulation of choline and purine nucleotide metabolites by SLC25A48. **A.** Schematic of pre-brown adipocytes isolated from SLC25A48-KO mouse to generate stable cell line with SLC25A48-KO and rescue of SLC25A48 (SLC25A48-KO^SLC25A48-Flag^). Pre-brown adipocytes were isolated from SLC25A48-KO brown adipose tissue and immortalized with SV40 through retroviral infection. Subsequently, expression of SLC25A48 was rescued in SLC25A48-KO brown adipocytes through retroviral expression of SLC25A48-Flag cDNA (SLC25A48-KO^SLC25A48-Flag^) to generate a rescue cell line from the same population of cells. **B.** Protein expression of SLC25A48 in SLC25A48-KO^SLC25A48-Flag^ brown adipocytes though immunoblotting for Flag. **C.** Western blot for mitochondrial complex proteins in brown adipocytes. Bands top to bottom complex IV (CV, ATP5A), complex III (CIII, UQCRC2), complex IV (MTCO1), complex II (SDHB), complex I (NDUFB8). N = 3 per group. **D.** Extracellular acidification rate (ECAR) of SLC25A48-KO and SLC25A48-KO^SLC25A48-Flag^ in response to norepinephrine (NE), oligomycin (oligo), carbonyl cyanide-p-trifluoromethoxyphenylhydrazone (FCCP), and rotenone and antimycin A (Rot/AA). N = 10 per group. Statistic is unpaired t-test. Circles represent mean and error shown as s.e.m.

**Supplemental Figure 3.**
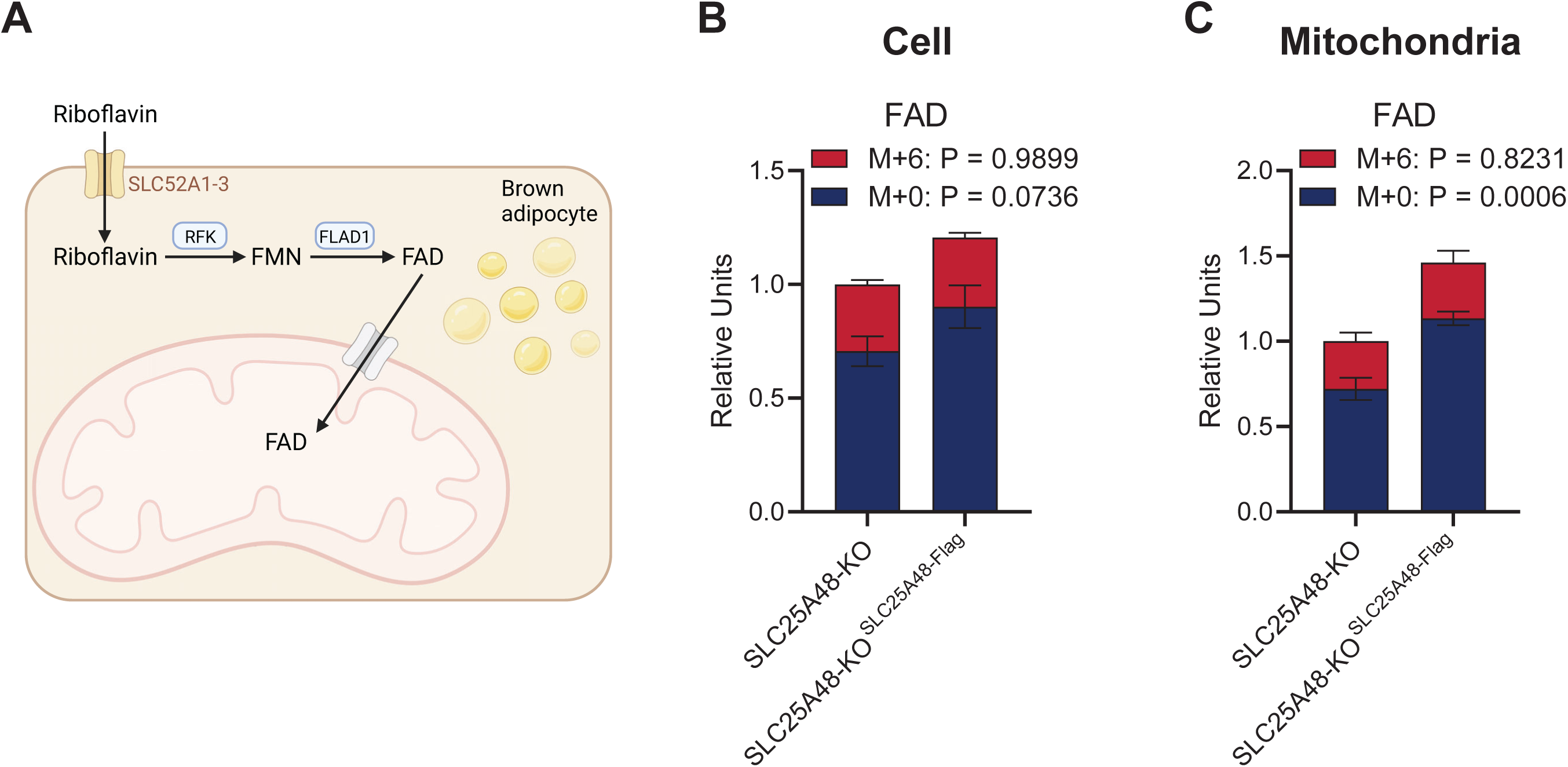
SLC25A48 does not mediate the import of FAD into mitochondria. **A.** Schematic of mitochondrial FAD transport tracing assay. Brown adipocytes were treated with labeled riboflavin (^13^C_4_,^15^N_2_ Riboflavin; M+6) for 4 hours. Riboflavin is imported into cells through plasma membrane transporters SLC52A1, SLC52A2, SLC52A3. In the cytosol, riboflavin is converted to flavin mononucleotide (FMN) by riboflavin kinase (RFK) and FMN is converted to flavin adenine dinucleotide (FAD) by flavin adenine dinucleotide synthetase (FLAD1). FAD import into the mitochondria is not fully understood. After 4 hours of treatment, cells were scraped and split into whole cell and mitochondrial fractions. **B.** Whole cell labeled (M+9, red) and unlabeled (D0, blue) FAD in SLC25A48-KO and SLC25A48-KO^SLC25A48-Flag^ brown adipocytes. Statistic: two-way ANOVA with Šídák’s multiple comparisons test. Values relative to total FAD (M0 and M+9) of Slc25a448-KO. Bars represent mean and error shown as s.e.m. **C.** Mitochondrial labeled (M+9, red) and unlabeled (D0, blue) FAD in SLC25A48-KO and SLC25A48-KO^SLC25A48-Flag^ brown adipocytes. Statistic: two-way ANOVA with Šídák’s multiple comparisons test. Values relative to total FAD (M0 and M+9) of SLC25A48-KO. Bars represent mean and error shown as s.e.m.

**Supplemental Figure 4.**
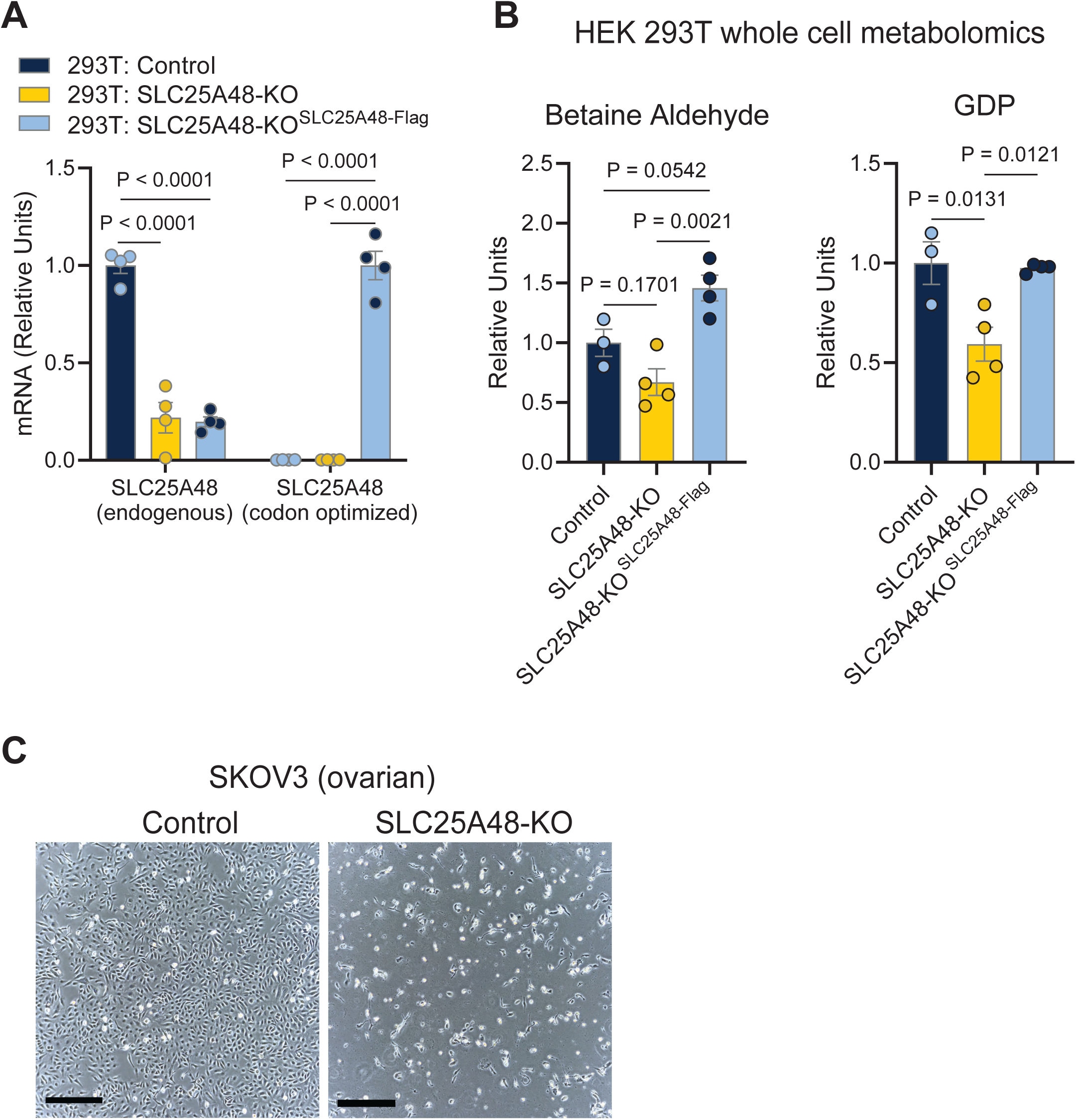
Mitochondrial choline catabolism via SLC25A48 regulates cell proliferation and survival. **A.** Expression of SLC25A48 (endogenous) and SLC25A48 (codon optimized) in HEK 293T control, SLC25A48-KO, and SLC25A48-KO treated with SLC25A48-Flag cDNA (SL25A48-KO^SLC25A48-Flag^). Endogenous expression relative to control and codon optimized relative to SLC25A48-KO^SLC25A48-Flag^. N = 4 per group. Statistic: two-way ANOVA with Tukey’s multiple comparisons test. Bars represent mean and error shown as s.e.m., individual values presented. **B.** Whole cell metabolite abundance of betaine aldehyde, guanosine diphosphate (GDP) of control, SLC25A48-KO, and SLC25A48-KO^SLC25A48-Flag^ HEK 293T cells. N = 3 per goup. Statistic: one-way ANOVA with Tukey’s multiple comparisons test. Bars represent mean and error shown as s.e.m., individual values presented. **C.** Representative image of SKOV3 ovarian cancer cells transfected with control or SLC25A48-KO plasmids for 24 hours. Scale bar: 500 μm.

